# Genetic editing of primary human dorsal root ganglion neurons using CRISPR-Cas9 with functional confirmation

**DOI:** 10.1101/2024.04.02.587857

**Authors:** Seph Palomino, Katherin Gabriel, Juliet Mwirigi, Anna Cervantes, Peter Horton, Geoffrey Funk, Aubin Moutal, Laurent Martin, Rajesh Khanna, Theodore Price, Amol Patwardhan

**Affiliations:** Department of Anesthesiology and Pain Management, University of Texas Southwestern Medical Center; 6202 Harry Hines Blvd., 9th Floor Dallas, Texas 75235; Department of Neuroscience and Center for Advanced Pain Studies, University of Texas at Dallas 800 W Campbell Rd, Richardson, TX 75080, USA; Southwest Transplant Alliance. 8190 Manderville Ln, Dallas, TX 75231, USA; Department of Pharmacology and Physiology, Saint Louis University,1402 S. Grand Blvd., St. Louis, Mo. 63104, USA; Department of Pharmacology, University of Arizona, 1501 N Campbell Ave, Tucson, AZ 85721, USA; Department of pharmacology and therapeutics, University of Florida, 1200 Newell Drive, ARB R5-234 Gainesville, FL 32610-0267

## Abstract

CRISPR-Cas9 editing is now the leading method for genome editing and is being advanced for the treatment of human disease. CRIPSR editing could have many applications for treatment of neurological diseases, including pain but traditional viral vector delivery approaches have neurotoxicity limiting their use. Overcoming these issues could open the door for genome editing treatments for diseases like intractable pain where the dorsal root ganglia (DRG) would be the desired target. To this end, we describe a simple method for viral-vector-independent transfection of primary human DRG (hDRG) neurons for CRISPR-Cas9 editing. As proof of principle, we edited *TRPV1, NTSR2*, and *CACNA1E* using a lipofection method with CRISPR-Cas9 plasmids containing reporter tags (GFP or mCherry). Transfection was successful as demonstrated by the expression of the reporters as early as two days *in vitro*. CRISPR-Cas9 editing was confirmed at the genome level with insertion and deletion detection system T7-endonuclease-I assay; protein level with immunocytochemistry and Western blot; and functional level through capsaicin-induced Ca^2+^ accumulation in a high-throughput compatible fluorescent imaging plate reader (FLIPR) system. This work establishes a reliable, target specific, non-viral CRISPR-Cas9-mediated genetic editing in primary human neurons with potential for future clinical application for intractable pain.

**Teaser:** We describe a non-viral transfection method for CRISPR-Cas9 gene editing in human dorsal root ganglion neurons.

## Introduction

Genome editing techniques have revolutionized the biomedical field, expanding our knowledge of human diseases and providing therapeutic applications in humans. Clustered regularly interspaced short palindromic repeats (CRISPR) - CRISPR associated protein 9 (Cas9) is a genome editing technique that uses a guide RNA (gRNA) to efficiently induce double stranded breaks in DNA, driving activation of DNA repair systems that can introduce site specific genomic modifications (*1, 2*). CRISPR-Cas9 systems have been widely utilized in understanding various disease related pathways and have accelerated application of genetically engineered cells and organisms. Previously, screening in both cell lines and primary cultures mostly involved small interfering RNA (siRNA), or RNAi, to investigate loss-of-function or drug activity of specific target proteins; However, the approach involves extensive troubleshooting to predict its partial knockdown effect, off-target effects, and is limited to protein-coding genes (*3, 4*). Cas9-mediated CRISPR tools are now widely used in screening due to their efficiency and ability to target the entire genome (*5, 6*). CRISPR-Cas9 has now entered therapeutic era with successful treatment of sickle cell disease and β-thalassemia (*7-10*). Despite these important advances, two of the biggest challenges encountered in genome editing are low transfection efficiency and toxicities arising from delivery strategies. These issues are particularly relevant for the nervous system because neurons are challenging to transfect without viral vectors and dorsal root ganglion toxicities from viral vectors have presented challenges in clinical trials and in preclinical development in non-human primate species (*11-15*).

There are several transduction/transfection methods available for CRISPR-Cas9 and they are divided into two broad categories: viral and non-viral (*16*). Viral transduction uses a viral vector to carry a specific DNA sequence into the host cells. For stable transductions retroviruses and lentiviruses are used, while adeno-associated viruses (AAVs) are used for transient transductions (*16*). The AAVs are widely recognized as highly effective at achieving transduction on notoriously difficult to transfect primary cells, but DRG toxicity is a clinically relevant concern of this approach (*11-15*). For non-viral transfection methods, there are multiple options such as mechanical, physical, or chemical transfections. Electroporation is the best-known physical method used to transfect cells and has been recently used to create the CRISPR-Cas9–edited CD34^+^ hematopoietic stem cells used in a sickle cell disease clinical trial (*10, 17*). Sonoporation, laser, or magnet -assisted transfection are also common methods but can irreversibly damage the cell membrane and have low transfection efficiency (*18*). Chemical transfections now mostly rely on lipid-based methods where positively charged lipid particles fuse with the phospholipid bilayer and enable the genetic material to easily enter the cell (*16*). Lipid-based nanoparticle approaches are used medically for vaccine delivery, gene therapy and are used in clinical trials for a variety of diseases (*19, 20*).

We sought to develop a viral-vector-independent method for CRISPR-Cas9 editing in human DRG neurons recovered from organ donors. Our work was motivated by the need for a genome editing technique for discovery work on these human neurons, and for potential clinical application where viral-vector-independent gene editing could have immediate impact such as for the treatment of intractable, inherited pain disorders like inherited erythromelalgia (*21*). To these ends, we describe a non-viral, lipofectamine-based transfection method for CRISPR-Cas9 genome editing in primary human DRG neurons recovered from organ donors. Successful transfection was verified by the expression of reporter protein and by mismatch insertion and deletion (indel) assay. We verified genetic editing at the protein level via immunocytochemistry and Western blotting demonstrating up to a 70% decrease in protein expression over a 5-day period after editing. Lastly, we demonstrate a functional loss of TRPV1 expression using a capsaicin-evoked Ca^+2^ imaging FLIPR assay. Our findings reveal a new method that can be used for assessment of gene function directly in human sensory neurons, for screening of drug action on human sensory neurons, and that can potentially be adapted for clinical use in diseases like intractable pain.

## RESULTS

### CRISPR-Cas9 transfection protocol and viability in human DRG cultures

Human DRG (hDRG) cultures were prepared through recovery of lumbar and thoracic DRGs from 12 organ donors (**Table 1**). The hDRGs were collected and immediately placed in cold (∼4°C) bubbled aCSF then transported to the laboratory for processing. Human DRG neuronal cultures were prepared as described previously and outlined in **Fig. 1** (*22*). We selected the transient receptor potential vanilloid 1 channel (*TRPV1*) as our first genetic target of interest due to its high expression in the hDRGs (*23*) and for its specific and thoroughly validated agonist, capsaicin (*24-26*). The CRISPR construct, designed by GeneCopoeia, contains a U6 promoter followed by a single guide RNA for the target of interest and the reporter tag gene for mCherry; The Cas9 nuclease gene is driven by a CMV promoter (**Fig. 2B**). All CRISPR constructs were obtained in a BSL-1 *E. coli* bacterial vector, the DNA plasmid was amplified through growth of the vector through ampicillin selective growth media and extracted using a maxi prep Qiagen kit.

**Table 1.** Human donor information. Donor information for all the human DRG samples used to conduct the experiments.

| Donor | Age | Sex | COD |
| --- | --- | --- | --- |
| 1 | 50 | Female | Anoxia |
| 2 | 20 | Male | Drug OD |
| 3 | 39 | Female | Stroke |
| 4 | 23 | Male | GSW |
| 5 | 34 | Male | MVA |
| 6 | 45 | Male | Anoxia |
| 7 | 39 | Female | Stroke |
| 8 | 22 | Male | MVA |
| 9 | 22 | Male | GSW |
| 10 | 37 | Male | MVA |
| 11 | 45 | Male | Head Trauma |
| 12 | 19 | Male | Anoxia |
| 13 | 38 | Male | Stroke |

GSW-Gunshot wound, MVA-Motor vehicle accident

**Figure 1.**
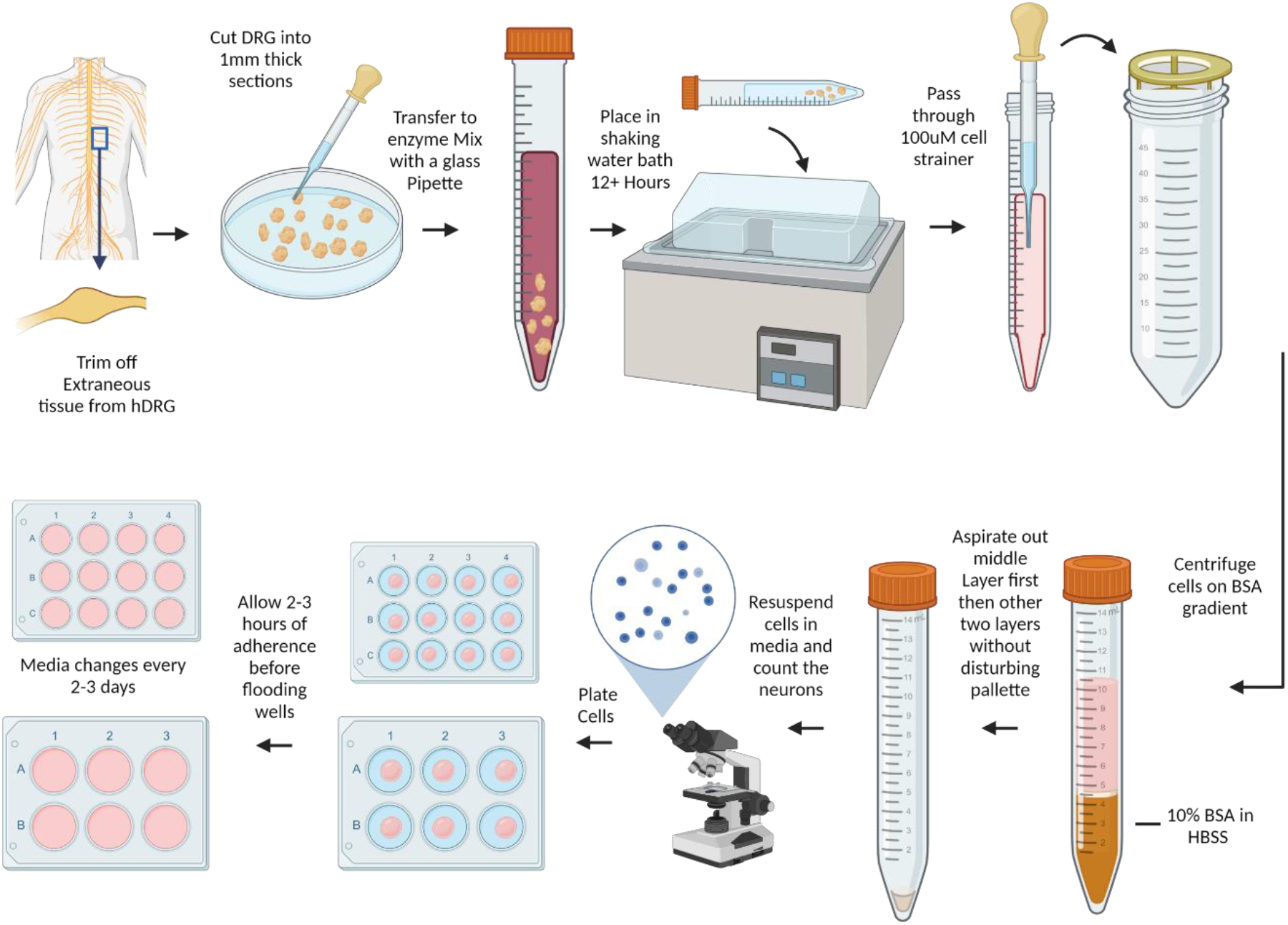
Human DRG dissociation protocol. The recovered DRG was trimmed to expose the cluster of cell bodies and minced into <3mm fragments. The tissue fragments were incubated in enzyme solution in a 10 mL conical tubes and placed in a 37°C shaking water bath for > 6 hours until the tissue completely dissociated resulting in a cloudy solution. The solution containing dissociated neurons was passed through a 100 μm sterile cell strainer and spun down in a 10% Bovine Serum Albumin/HBSS gradient. The resulting supernatant was removed and the palleted cells were resuspended in pre-warmed hDRG media. The cells suspension was carefully plated onto the appropriate dish and allowed to adhere for 3 hours at 37°C and 5% CO2 The wells were flooded with prewarmed complete DRG media and the cultures were maintained for up to 15 days with media changes were performed every other day (*22*).

**Figure 2.**
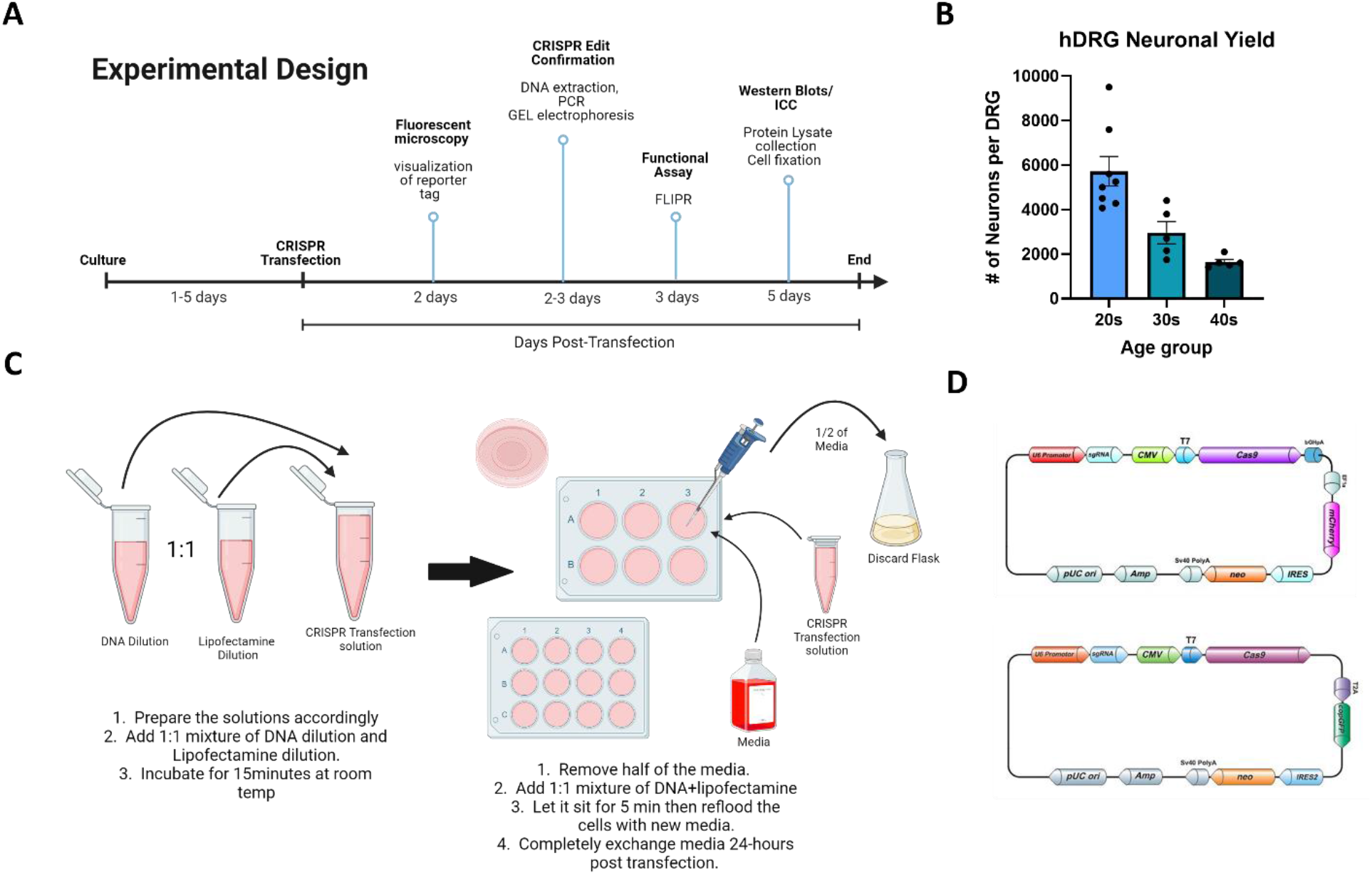
CRISPR-Cas9 Transfection experimental design and Protocol. **A**. General experimental timeline of CRISPR-Cas9 transfection in hDRG culture and each consequent experimental verification steps **B**. Measurement of neuronal yield per one hDRG dissociated across 3 different age groups. Each dot represents the neuronal yield for a single donor. **C**. Visual representation of transfection reagent and CRISPR plasmid formulation; a 1:1 mixture of DNA dilution and lipofectamine dilution incubated for 15 min before added to the hDRG cultured cells and incubated for 24-hours. **D**. Plasmid maps for CRISPR-Cas9 gene editing, one containing an mCherry reported tag and another containing a GFP reporter tag. Plasmid map images were obtained from GeneCopoeia website.

Our group’s experimental timeline for the various methods used for confirmation of the CRISPR genetic editing and functional testing is outlined in **Fig. 2A**. We chose a cationic lipid-based transfection method, using lipofectamine3000, to insert the plasmids into the cells present in the hDRG cultures. The concentration of the plasmid DNA used for the DNA-lipid complex ranged from 0.5-5μg depending on the size of the culture dishes and relative cell number. The amount of lipofectamine3000 was adjusted according to media volume and surface area of the culture dishes; the formulation of the plasmid-transfection reagent was adjusted according to the manufacturer’s recommendations and diluted in serum free BrainPhys culture media (**Fig. 2C) (Table 2)**. The extracted CRISPR-Cas9 plasmid was mixed with the P3000 reagent (2uL/ug DNA) and diluted in serum free BrainPhys culture media. The media containing lipofecitimine3000 and media containing the DNA-p3000 complex were mixed at a 1:1 ratio and incubated for 15 minutes at room temperature. At least half of the media was removed from the wells before the DNA-lipid complex was added and incubated for 5 minutes at room temperature. The wells were then refilled with fresh media and left for 24 hours in the incubator before a media change.

**Table 2.** Lipofectamine 3000 reagent protocol. The hDRG cells were transfected according to these optimized specifications.

| Component | 96-Well Plate | 12-Well Plate | 6-Well Plate |  |
| --- | --- | --- | --- | --- |
| BrainPhys Media | 4.85uL | 58.5uL | 121.5uL | Dilute lipofectamine in media and mix well. |
| Lipofectamine 3000 | 0.15uL | 1.5uL | 3.75uL |  |
| BrainPhys Media | Adj to 5uL | Adj to 60uL | Adj to 125uL | Dilute DNA in media and add p3000 Accordingly. |
| DNA* | 0.5-1ug | 1.5-3ug | 3-5ug |  |
| P3000 Reagent | 2uL/ug of DNA | 2uL/ug of DNA | 2uL/ug of DNA |  |
| Diluted DNA | 5uL | 60uL | 125uL | Add 1:1 mixture and incubate for 15min |
| Diluted Lipofectamine | 5uL | 60uL | 125uL |  |

We compared the overall neuronal yield from each hDRG cultured across different age groups and determined that older donors produce an overall lower neuronal yield (**Fig. 2B**). This information led us to determine an age limit of 50 years old to maximize yield and survivability of neurons in culture. Next, we determine if the health of the culture was affected by the number of days the cells were exposed to the plasmid/lipid complex by performing a cell viability MTS assay. This assay enabled us to calculate the change in viability of the cells before and after CRISPR-Cas9 and/or cationic lipid treatment with either 24- or 48-hour exposure time (**Fig. 3A**). Although the change in viability was not significant there was a clear negative trend associated with longer exposure time. This led us to determine that the exposure of the lipid-plasmid complex should not extend beyond 24 hours.

**Figure 3.**
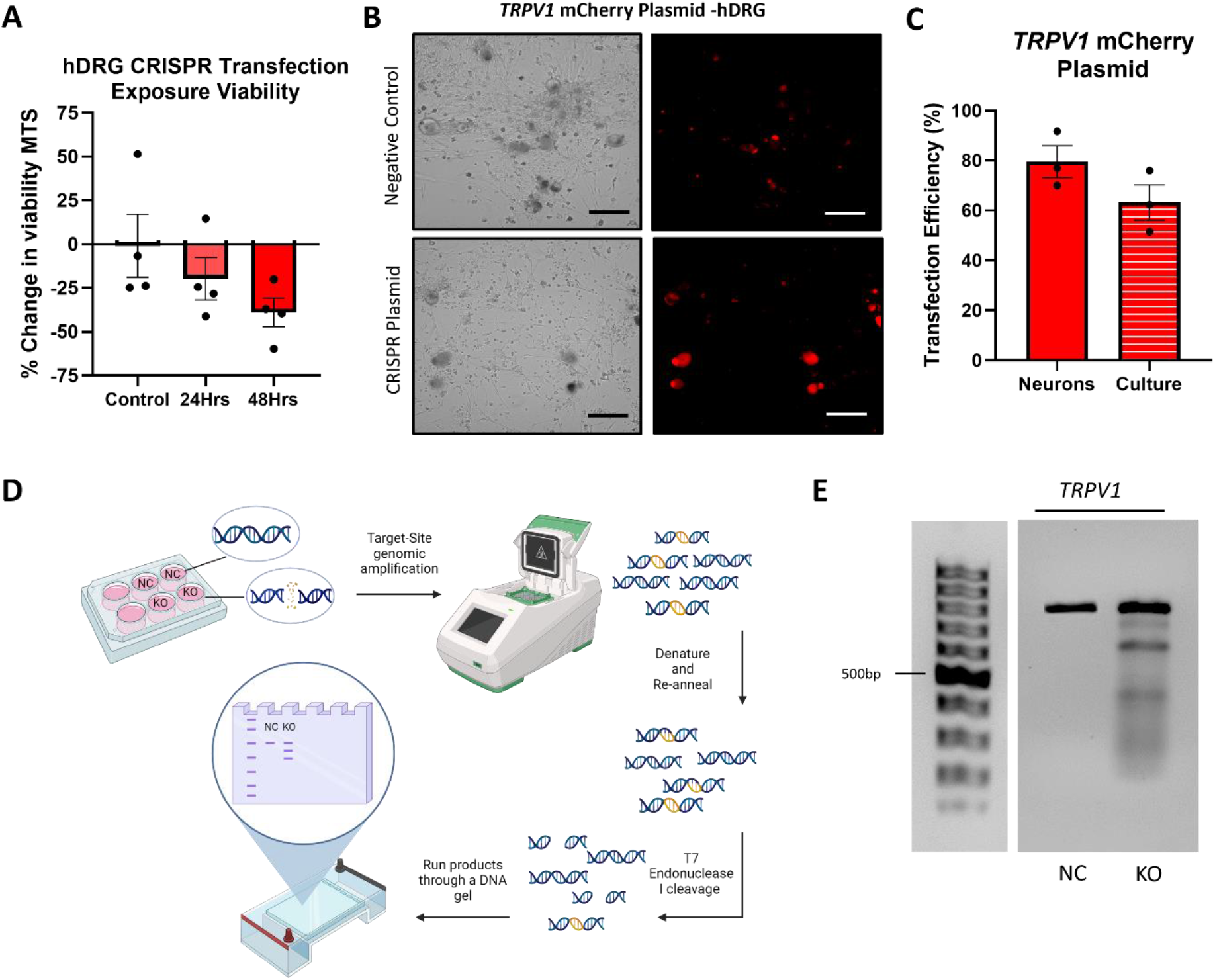
Validation of CRISPR-Cas9 transfection with reporter tag and T7 endonuclease assay. **A**. MTS % change in viability in hDRG cultured cells treated for either 24- or 48 - hours with the *TRPV1* gRNA plasmid (500ng) demonstrating cells should not be exposed to the lipid-plasmid complex for more than 24-hours. **B**. Immunofluorescent images of mCherry reporter tag expression in the CRISPR-Cas9 *TRPV1* KO plasmid treated cells (bottom), compared to the negative control cells (top). Representative brightfield images (left) and the fluorescent images (right) were taken using a Biorad ZOE Fluorescent cell imager. Scale bar 100μm **C**. % transfection efficiency for the *TRPV1* CRISPR plasmid quantified by fluorescent tag expression in neurons alone and in the entire hDRG culture. Each point represents the average of at least 100 neuronal and at least 600 non-neuronal cells quantified from 3 individual replicates. The transfection efficiency for the culture includes both neuronal and non-neuronal cells. **D**. Visual workflow of T7 endonuclease I assay used to confirm CRISPR-Cas9 genetic edit. **E**. Representative image of DNA gel demonstrating multiple DNA band fragments in the *TRPV1* CRISPR Plasmid exposed sample compared to the single negative control sample band.

### Validation of CRISPR-Cas9 transfection with reporter tag and T7 endonuclease assay

To validate the plasmid transfection of hDRG neurons, we looked for expression of the mCherry reporter tag in the DRG cultures exposed to the DNA-lipid complex and compared them to cultures that received lipofectamine without plasmid DNA (negative control). A common characteristic of primary neuronal cultures is the presence of autofluorescence due to lipofuscin, a lipid containing pigment that accumulates in the cytoplasm due to aging (*27*). The lipofuscin can be seen in the bright field images (left) as the darker pigmented areas in the neurons, and the corresponding autofluorescence in the fluorescent images (right) (**Fig. 3B)**. Aside from the autofluorescence from the lipofuscin, compared to the negative CRISPR control, the *TRPV1* gRNA transfected cells showed a robust expression of the mCherry tag showing greater than 60% transfection efficiency (**Fig. 3B&C**). Since the primary cultures are a mixed culture, we quantified the expression of the reporter tag to determine the transfection efficiency in neurons alone, and in the entire culture including neurons (culture) (**Fig 3C**). We observed the tag beginning to express in culture, from all donors of any age, two days after transfection. These findings demonstrate that a lipid transfection method was successful in delivering DNA plasmid into cells of a primary hDRG culture.

The CRISPR-Cas9 double stranded DNA damage can be recognized and repaired by non-homologous end-joining (NHEJ) and cause frameshift mutations, premature stop codons, or damage to the open reading frame of target genes by insertions or deletions (indels) (*28*). These indel errors are exploited to verify target site editing by using mismatch detection assays. Since the reporter tags began to express two days post-transfection, we chose the same timing to extract DNA to verify editing. For this verification step we chose to use a T7 endonuclease I enzyme which recognizes and cleaves mismatched DNA. Post DNA extraction, we generated PCR products using primers flanking the targeted gene sequences, then used heat denaturation followed by a room temperature re-annealing step allowing them to generate double stranded fragments that contain mismatched DNA (**Fig. 3D**). The T7 endonuclease I enzyme can detect and cleave mismatched DNA which can be visualized in an agarose gel. The enzyme cleaved the DNA of the cells exposed to the *TRPV1* CRISPR plasmid but not the negative control cell’s DNA as demonstrated by the strand breaks in the DNA gel (**Fig. 3E**). This confirmed successful DNA editing of the *TRPV1* gene in CRISPR treated hDRG cultures.

### Changes in protein expression of CRISPR-Cas9 treated hDRG neurons

To further validate specific editing of the target gene, *TRPV1*, we performed immunocytochemistry (ICC) and western blotting to assess changes in protein expression in the hDRG cultures. Five days post transfection, we fixed cells on cover slips for ICC and collected hDRG whole cell lysates from 6-well plates for Western Blot quantification of TRPV1 protein expression. We chose peripherin, an intermediate filament enriched in peripheral neurons (*29*) and expressed by all hDRG neurons (*30*), to label neurons in ICC coverslips. The respective reporter tags associated with the CRISPR plasmids were still expressed five days post transfection (**Fig. 4A**). Moreover, when compared to the negative control cells, TRPV1 expression was significantly decreased; Showing a 70% reduction in fluorescent intensity of TRPV1 protein expression in neurons labeled by peripherin (**Fig. 4B**). We quantified the total change ofTRPV1 expression in the hDRG culture by Western Blot from the supernatant of the collected whole cell lysate. The Western blot demonstrates a statistically significant 50% decrease in TRPV1 expression (**Fig. 4C**). Overall, this data demonstrates successful editing and reduced TRPV1 protein expression in hDRG culture.

**Figure 4.**
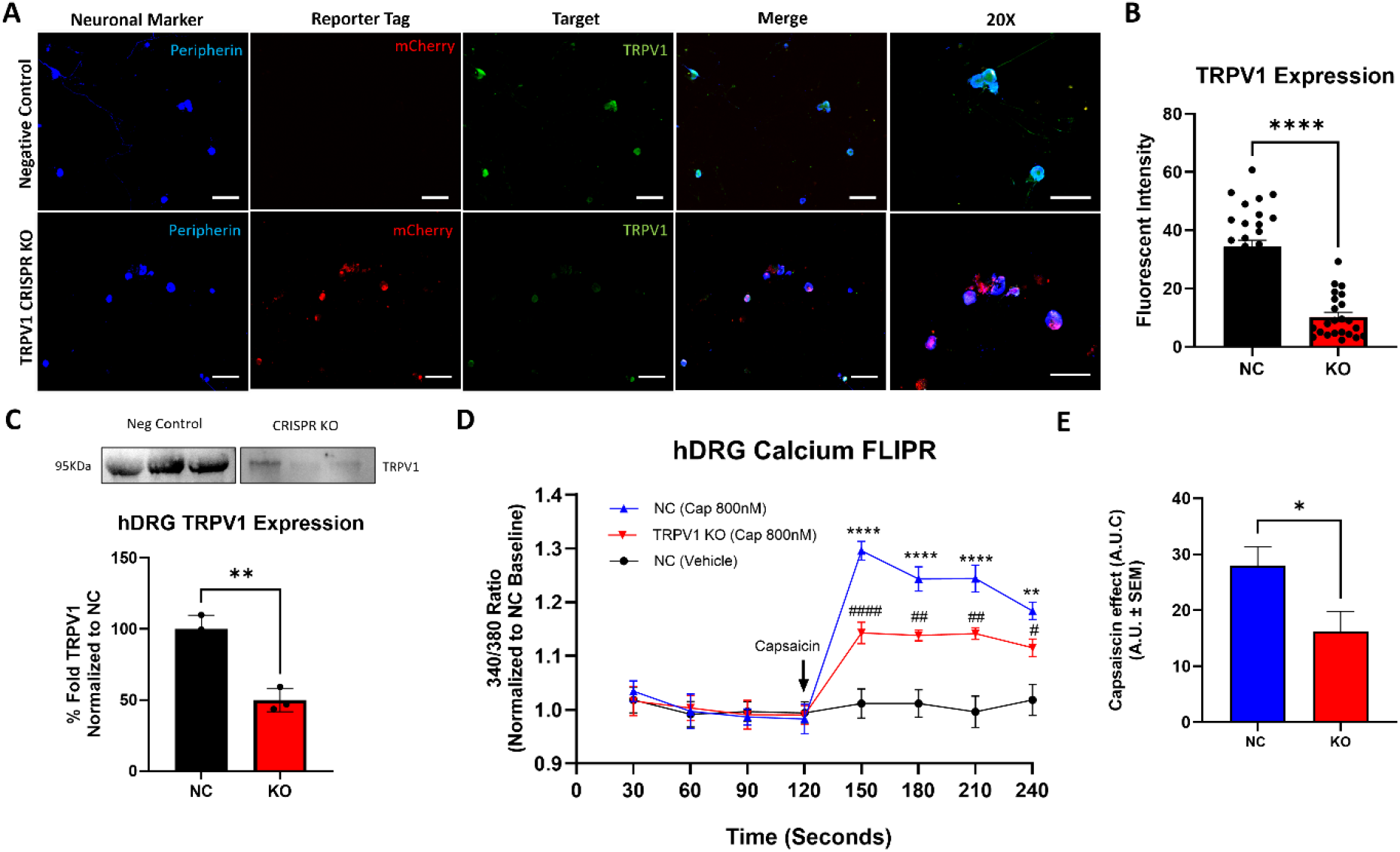
Validation of decreased protein expression and protein function in *TRPV1* CRISPR plasmid treated cells. hDRG cultures were transfected with CRISPR-Cas9 plasmid targeting *TRPV1* and the cells were either fixed or protein lysates were collected 5 days later and analyzed via western blots or immunocytochemistry. **A**. Representative 10X fluorescent images staining for TRPV1 in negative control or CRISPR-Cas9 treated hDRG cultures. Peripherin (blue) was used as the neuronal marker, the reporter tag (mCherry) was seen only in the CRISPR treated cells, while the target of interest is significantly decreased in the CRISPR treated cells. The merge images of all three channels are shown for each negative control and CRISPR treated cells with the corresponding 20X image. **B**. Data for the mean fluorescent intensity of TRPV1 (Green) was quantified as means ± SEM of n=10 neurons per group in 3 individual replicates. ****P < 0.0001 by unpaired two-tailed t test. Scale bars = 100μm **C**. The representative Western blot and quantification of TRPV1; data was normalized to total protein, then further normalized to the negative control group (100%). ** = p < 0.01 by unpaired 2-tailed t test. **D**. hDRG cultures were transfected with CRISPR-Cas9 plasmid targeting *TRPV1*. 72-hours post-transfection, the cells were incubated with Tyrode’s buffer containing 2 uM Fura 2-AM/ Pluronic F-127 covered at room temperature for an hour then incubated with Tyrode’s buffer supplemented with 2.5 mM probenecid for 20-30 minutes. A 2-minute baseline was recorded followed by 800nM capsaicin stimulation recorded for another 2-minutes in 30 second timepoint intervals. A decreased capsaicin evoked calcium response is seen in the TRPV1-CRISPR-KO cells compared to the NC capsaicin stimulated cells. The data was reported as the 340/380 ratio and normalized to a 4-point negative control baseline. NC-Capsaicin vs NC vehicle same time point; **** = p < 0.0001; ** = p < 0.01; * = p < 0.1; and NC-Capsaicin vs TRPV1-KO-Capsaicin same timepoint; #### = p < 0.0001; ## = p < 0.01; # = p < 0.1; 2-Way ANOVA with Tukey’s post hoc test. Data was measured in two separate donors with 4-6 individual replicates per group. **E**. Representative area under the curve (AUC) comparing NC capsaicin response to TRPV1-CRISPR KO capsaicin response post 120 second timepoint. * = p < 0.05 by unpaired 2-tailed t - test.

### TRPV1 functional expression is decreased after CRISPR-Cas9 knockout in hDRG neurons

To demonstrate a decrease in protein function in response to the TRPV1 knockout we utilized a fluorescent imaging plate reader to detect changes in intracellular Ca^2+^ levels. Using Fura 2 AM dye we quantified the 340/380 ratio in response to capsaicin, an established specific TRPV1 agonist or vehicle. The cells were seeded into a 96-well plate and transfected 24-hours later with 500 ng of CRISPR plasmid plus lipofectamine. Negative control wells received only lipofectamine treatment with the same timeline as the DNA-lipid complex treated wells. Three days post transfection the cells were incubated with Fura-2 AM/Pluronic F-127 (1:1) in Tyrode’s buffer for one hour at room temperature, followed by 30-minute incubation with Tyrode’s with 2.5 mM probenecid to improve intracellular indicator retention. To assess functional knockout of *TRPV1*, we recorded a 2-minute baseline followed and proceeded to stimulate the cells with 800 nM capsaicin. We recorded the fluorescent intensity for 2-minutes post stimulation in 30-second intervals. Compared to the negative control wells, the *TRPV1*-edited wells had a significantly reduced response to 800nM capsaicin; The cells exposed only to vehicle showed no change in their intracellular Ca^2+^ concentration (**Fig. 4D**). We conclude CRISPR-Cas9 editing of the *TRPV1* gene in hDRG culture results in decrease of functional receptor activity 3 days post transfection.

### CRISPR-Cas9 genetic editing of *NTSR2* and *CACNA1E* in hDRG cultures

We chose two other targets for editing, the G-protein coupled receptor (GPCR) neurotensin receptor 2 *(NTSR2*), and the R-type voltage gated calcium channel Cav2.3 (*CACNA1E*). These plasmids were also designed to contain a U6 promoter followed by a single guide RNA for the target of interest with the Cas9 being driven by a CMV promoter, but instead of mCherry, we chose to try a GFP reporter tag (**Fig. 2D**). Like the mCherry plasmid, the expression of the GFP tag began 2 days post transfection and both *CACNA1E* and *NTSR2* gRNA transfected cells showed robust expression of the GFP reporter tag compared to the negative control cells; showing an approximate 50% and 60% respective transfection efficiency (**Fig. 5A-C**). To determine if there were differences in viability between the cells treated with the three different CRISPR plasmids or their respective tags, we performed an ATP luminescent assay. Compared to the control non-CRISPR-treated cells, the cells exposed to the GFP lipid-plasmids displayed a statistically significant decrease in viability, but there were no statistically significant differences between the control cells and the *TRPV1* CRISPR plasmid transfected cell (**Fig. 5D**). It is important to note, there were no statistically significant differences between the control cells and the cells treated with Lipofectmaine3000 alone. Nor was there a significant difference between the Lipofectamine3000 treatment compared to the DNA-lipid complex treated cultures. Overall, the cells exposed to the plasmid containing the mCherry tag were easier to confirm transfection as we could better differentiate between the lipofuscin and the fluorescent tag expression (**Fig. 3B**). This led us to conclude that the plasmid containing the mCherry reporter tag may be preferential for future experiments to ensure better visualization of expression.

**Figure 5.**
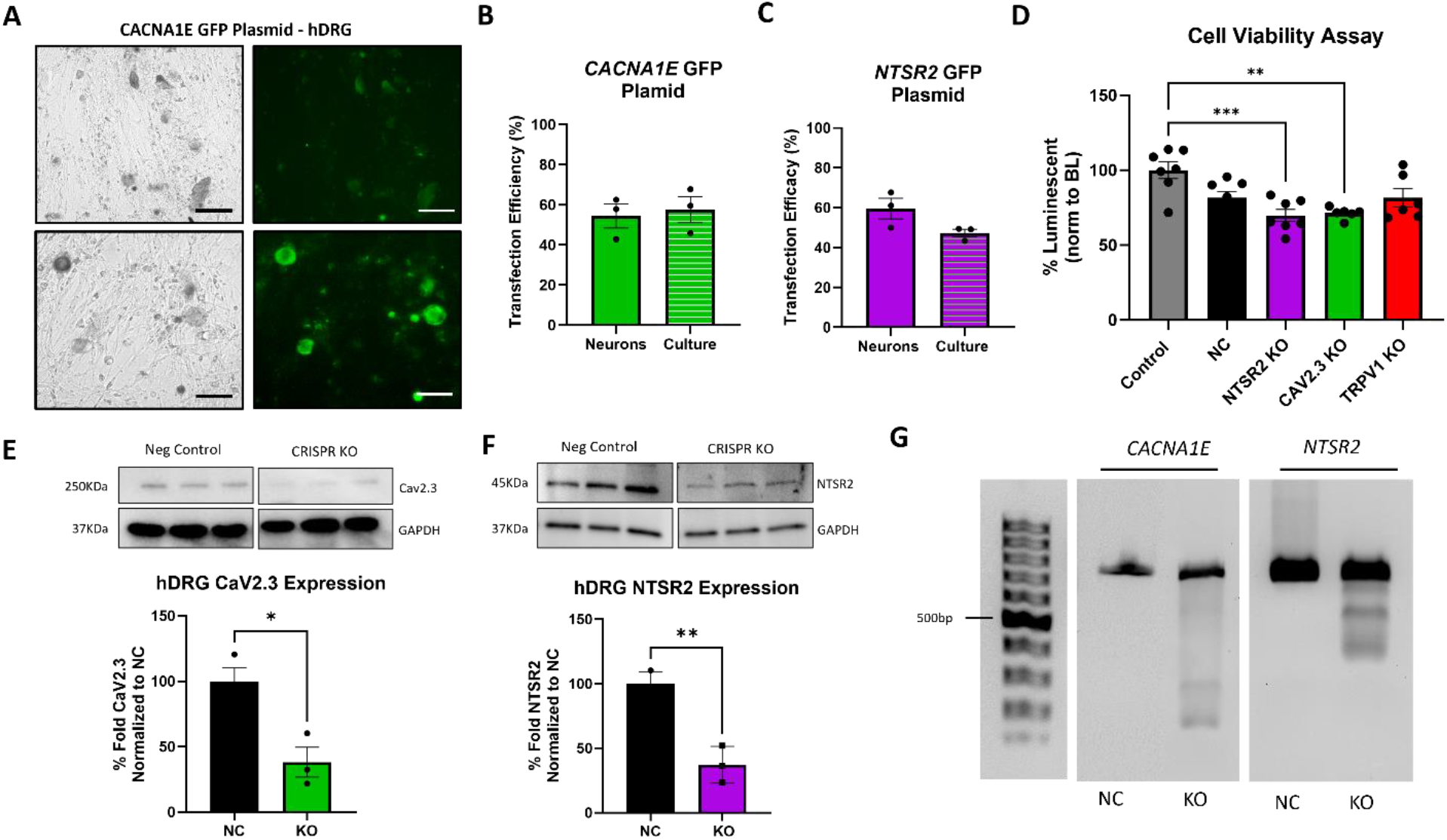
CRISPR-Cas9 genetic editing of *NTSR2* and *CACNA1E* in hDRG cultures. **A**. Representative immunofluorescent images of GFP reporter tag expression in the CRISPR-Cas9 *CACNA1E* KO plasmid treated cells (bottom), compared to the negative control cells (top). Representative brightfield images (left) and the fluorescent images (right) were taken using a Biorad ZOE Fluorescent cell imager. Scale bar 100μm. **B**. Percent transfection efficiency for the *CACNA1E* CRISPR plasmid quantified by fluorescent tag expression in neurons alone and in the entire hDRG culture. **C**. Percent transfection efficiency for the *NTSR2* CRISPR plasmid quantified by fluorescent tag expression in neurons alone and in the entire hDRG culture. (Images not shown). **D**. CellTiter Glo 2.0 Assay demonstrating viability of cells may differ between the cells exposed to plasmids containing the GFP compared to the mCherry reporter. Both *NTSR2* and *CACNA1E* DNA plasmids with GFP reporter tags show a significant decrease in viability compared to non-treated control cells but not in comparison to lipofectamine treated NC cells. Both lipofectamine alone and the plasmid containing the mCherry tag (*TRPV1*) show no significant decrease in viability compared to the non-treated control cells. Luminescence was measured across two different donors with 3 individual replicates per treatment group using a Tecan Spark 20 plate reader; *** = p < 0.001; ** = p < 0.01; by 1-Way ANOVA with Tukey’s post hoc test **E**. Representative Western blot and quantification of NTSR2; data was normalized to GAPDH, then further normalized to the negative control group (100%). ** = p < 0.01; * = p < 0.1 by unpaired 2-tailed t - test. **F**. Representative Western blot and quantification of Cav2.3; data was normalized to GAPDH, then further normalized to the negative control group (100%). * = p < 0.05 by unpaired 2-tailed t -test. **G**. Representative image of DNA gel demonstrating multiple DNA band fragments in the *NTSR2 & CACNA1E* CRISPR plasmid exposed sample compared to the single negative control sample band.

We validated the genetic editing of these targets (*NTSR2 & CACNA1E)* using the T7 endonuclease method as with the *TRPV1* plasmid in culture (**Fig. 5G**). We observed a significant reduction in protein expression via Western blot with a 62% reduction in Cav2.3 and a 63% reduction in NTSR2 expression in culture (**Fig. 5E&F)**. There was also a significant reduction of fluorescent intensity representative of Cav2.3 and NTSR2 in neurons expressing the reporter tag observed in ICC (**Fig. 6**). Therefore, we have validated the transfection method for two additional targets: one a GPCR and the other a voltage-gated ion channel.

**Figure 6.**
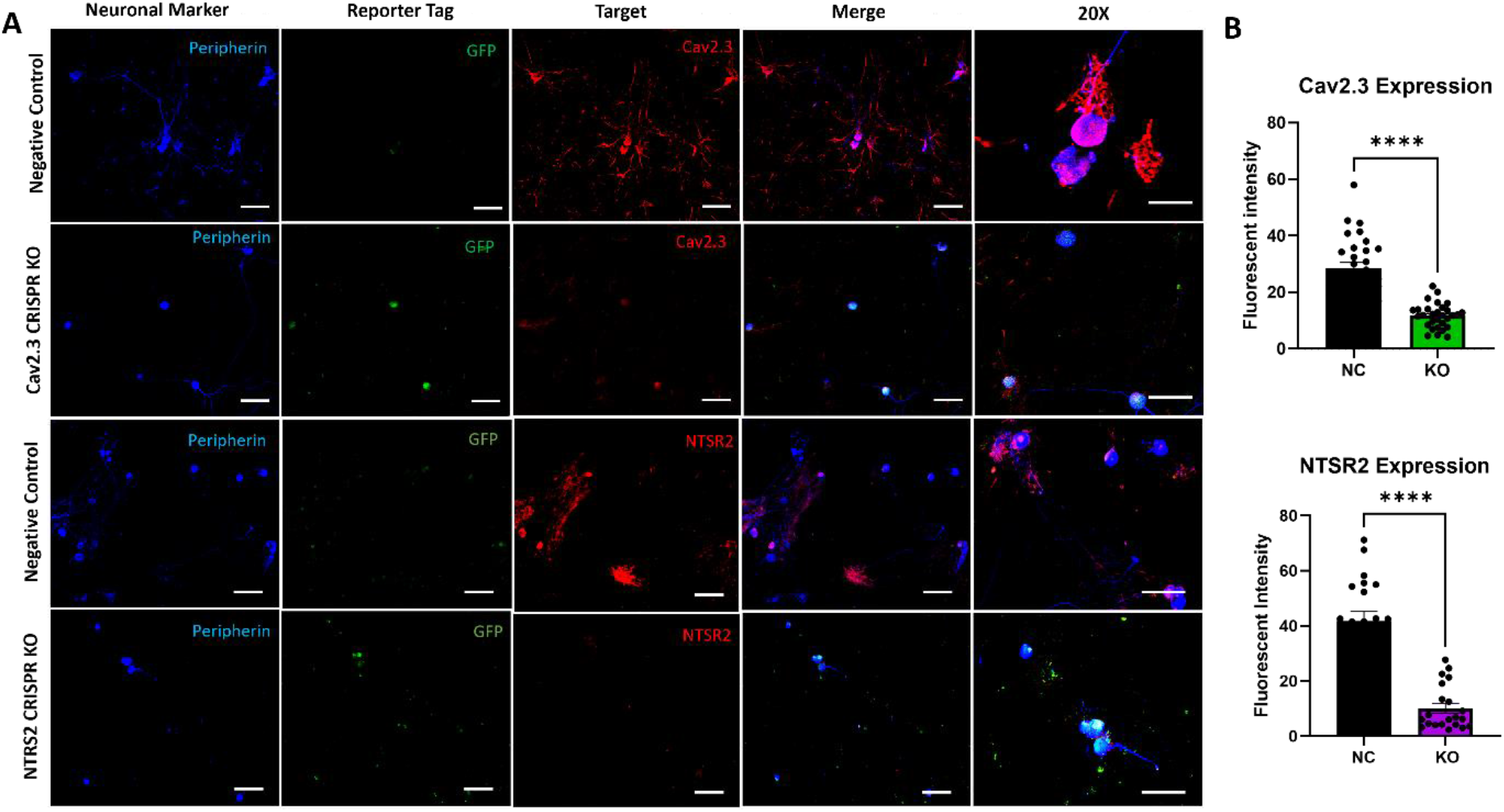
Immunocytochemistry validation of CRISPR editing of NTSR2 and Cav2.3 protein levels in hDRG culture. Representative 10X fluorescent images staining for NTSR2 and Cav2.3 of negative control or CRISPR-Cas9 treated hDRG cultures. **A**. Peripherin (blue) was used as the neuronal marker, the reporter tag (GFP) was seen only in the CRISPR treated cells, while the target of interest is significantly decreased in the perspective CRISPR treated cells. The merge images of all three channels are shown for each negative control and CRISPR treated cells with the corresponding 20X image. **B**. Data for the mean fluorescent intensity for the targets of interest Cav2.3 (Red), and NTSR2 (Red) was quantified as means ± SEM of n=10 neurons per group in 3 individual replicates. ****P < 0.0001 by unpaired two-tailed t-test. Scale bars = 100μm

## Discussion

In summary, we provide a non-viral lipid-based transfection method to perform CRISPR-Cas9 genomic edits in primary human DRG culture. Primary neuronal cultures are difficult to maintain, and several factors such as culturing conditions, donor age, and postmortem interval can affect the overall health of the culture (*31*). The primary culture method developed by our group was described in **Figure 1** and is a key component to maintaining a healthy DRG neuronal culture. The postmortem interval for all the donors was less than four hours, which is another key factor influencing culture viability and high cell yield. We also observed hDRG cultures from older aged donors typically showed lower neuronal cell yields and subsequently lower viability when subjected to lipofectamine transfection leading to cell loss in an already depleted neuronal population. For most of the experiments we preferentially used DRGs recovered from donors not exceeding 50 years of age (**Fig. 2B)** (**Table 1**).

The transfection efficiency can depend on the promoters used, so we chose plasmids containing the universal U6 promoter driving the sgRNA sequence, and the non-specific CMV promoter driving the Cas9 protein (**Fig. 2D**) (*32*). The plasmids also contained a reporter tag (GFP or mCherry) and a neomycin resistance gene for mammalian cell selection. Our initial step to verify if we achieved transfection was fluorescent imaging to observe the expression of the reporter tags (GFP or mCherry). The robust expression of the reporter tags in neurons demonstrated the lipofectamine is an efficient delivery method, and the promoters are sufficient at driving expression to cause genetic editing. Out of the three different CRISPR plasmid constructs tested, the minimum transfection efficiency was 50% for the *CACNA1E* construct (**Fig. 5B**). It is important to note, the lipid-plasmid complex should not be left in the culture media for more than 24 hours. This is in part due to the change in viability, but also because plasmids can lower the pH of the culture medium, and this can in turn affect the fluorescent intensity of the reporter tags (**Fig. 3A**)(*33*). It is possible to design the plasmid to contain a cell specific promoter to drive the Cas9 protein, and therefore lead to a targeted knockout in a cell specific manner (*34*). A widespread practice to identify and obtain an enriched transfected cell population is to FACS sort the cells using the reporter tag. In our case, due to the large diameter size and the fragility of the hDRG neurons, this was not a practical option. In addition, we chose not to use neomycin to preferentially select transfected cells as it is evident it can be toxic to mammalian cells, and we did not want to further risk the viability of hDRG cultures (*35*).

A CRISPR genomic edit can be confirmed using Sanger DNA sequencing or tracking of indels by decomposition (TIDE) assay, however we chose a simpler more time efficient method by using an indel assay with enzymatic digestion that can be performed 36-48 hours post transfection (**Fig. 3D**). This assay confirmed genetic editing in culture, specifically in the targeted regions of interest (**Fig. 3&5**). Confirmation of significant protein changes in culture were seen via western blots with at least a 50% (TRPV1) and more specifically in the neuronal population via immunocytochemistry with at least a 60% (Cav2.3) reduction in fluorescent intensity representative of protein expression across all three targets (**Fig. 4-6**). Taken together, we conclude achievement of high delivery efficiency of the CRISPR plasmids for our three targets of interest; Cav2.3, NTSR2, & TRPV1.

To test the effect of the knockout on protein function, we developed a protocol for measuring intracellular Ca^+2^ by using a fluorescent imaging plate reader (FLIPR) assay with the calcium indicator Fura 2AM dye in primary hDRG culture, a method which can be used in a high through put screening (HTS) fashion (**Fig. 4**) (*36, 37*). A FLIPR is a very efficient tool at detecting comprehensive and homogenous intracellular Ca^+2^ changes and has shown promise as a functional output in human iPSC-derived nociceptors (hiPSCdNC) (*38, 39*). TRPV1 edited wells showed a significant decrease in response to capsaicin compared to negative control wells; However, it did not abolish it completely as compared to non-stimulated cells (**Fig. 4**). This could be due to residual TRPV1 channel protein reserve present in the cells, as the experiment was performed on the 3^rd^ day post transfection. If we maintained the cells on these plates 5 days post transfection, as in the case of the coverslips and 6-well plates used in protein quantification, we may have seen a greater decrease in functional TRPV1. However, the small surface area in the 96-well plate becomes a limiting factor for how many days the cells can be maintained in culture before overgrowth of glial cells, overcrowding the neurons and the cells begin to detach. Some compromises between the type of primary culture and the degree to which editing will be reflected in protein turnover need to be considered in experimental design.

The advancement of nano-delivery systems has been beneficial in providing easy to use, reliable transfection of nucleic acids into target cells (*40, 41*). With the growing application of genomic engineering in the clinic, cationic lipid nanoparticles are more likely to have a greater impact over viral vectors. To our knowledge, we demonstrated the first successful lipid-based delivery method of CRISPR-Cas9 genome editing system in primary human DRG neurons. Overall, our work provides evidence that a cationic lipid delivery system is a viable transfection method for gene editing in primary human neurons which can be utilized in many cell-based assays. In summary, we provide a straightforward method for genome editing in human DRG neurons which will enable dissection of mechanisms and targets in these cells with precision and enables future work for therapeutic application.

## Materials and Methods

### Human DRG collection

All human tissue procurement procedures were approved by the Institutional Review Boards at the University of Texas at Dallas. Human DRGs were surgically extracted using a ventral approach (*42*) from organ donors within 4 hours of cross-clamp and placed immediately on artificial cerebrospinal fluid (aCSF). All tissues were recovered in the Dallas area via a collaboration with the Southwest Transplant Alliance. Tissue samples from 12 donors were used in this study, information is provided in Table 1.

### Human DRG dissociation protocol

The recovered DRG was trimmed with No. 5 forceps (Fine Science Tools, Cat#11252-00) and Bonn scissors (Fine Science Tools, Cat# 14184-09) to expose the cluster of cell bodies. At the time of culturing, a final enzyme solution consisting of 2mg/mL Stemzyme I (Worthington Biochemical, Cat#LS004106) and 0.1 mg/mL DNAse I was prepared in HBSS (Thermo Scientific, Cat#14170161) and allowed to incubate in a water bath. A total of 5 mL of enzyme solution was used to dissociate the hDRG tissue. Ganglia were finely minced with scissors into smaller fragments less than 3 mm (about 0.12 in). The tissue fragments were transferred to the prewarmed enzyme solutions in 10 mL conical tubes and placed in a 37°C shaking water bath until the tissue completely dissociated resulting in a cloudy solution (6-18 hrs in total). The solution containing dissociated neurons was passed through a 100 μm sterile cell strainer (VWR, Cat# 21008-950) into a 50 mL conical tube. The cell suspension was then gently added to a 3 mL 10% Bovine Serum Albumin (Biopharm, Cat#71040)/HBSS gradient and spun down at 900*xg* for 5 min at RTP, 9 acceleration, 5 deceleration. The resulting pallet of cells at the bottom of the tube was resuspended in prewarmed and sterile filtered DRG media. 20μL of cell suspension was carefully plated onto the 96-well plate, 120μL onto each coverslip, and 500μL onto the 6 well plates at a density of at least 150, 300, and 800 neurons respectively, and allowed to adhere for 3 hours at 37°C and 5% CO2. The wells were flooded with the appropriate amount depending on the plate of prewarmed complete hDRG media. Media changes were performed every other day. Primary hDRG neurons are non-dividing postmitotic cells that are difficult to maintain; therefore, to achieve the most optimal cell count and cell health for the transfection experiments, donors above 50 years old were excluded.

### Antibodies

Antibodies used were as follows: Peripherin (chicken 1:1000; EnCor, AB_2284443) Cav2.3 (Rabbit, Alomone labs, 1:100 AB_2039777; lot ACC006AN0302), NTSR2 (rabbit, 1:200, ThermoFisher BS-12004R), TRPV1 (Rabbit, 1:500 Fisher scientific PA1748) GAPDH (mouse, Cell signaling 97166s) secondary Alexa Fluor goat anti-mouse immunoglobulin G (IgG) 594 (1:5000; Invitrogen, A11032, lot 1985396), and secondary Alexa Fluor goat anti-mouse IgG 488 (1:5000; Invitrogen A-11034, lot 2110499). Alexa Fluor goat anti-chicken IgG 647 (1:5000; Invitrogen A-21449, lot 2186435).

### Plating Media

BrainPhys® media (Stemcell technologies, cat. no. 05790), 1% penicillin/streptomycin (Thermo Fisher Scientific, Cat# 15070063), 1% GlutaMAX® (United States Biological, cat. no. 235242), 2% NeuroCult™ SM1 (Stemcell technologies, cat. no. 05711), 1% N-2 Supplement (Thermo Scientific, cat. no. 17502048), 2% HyClone™ Fetal Bovine Serum (ThermoFisher Scientific SH3008803IR), 1: 1000 FrdU, 10 ng/mL Human β nerve growth factor - (Cell Signaling Technology, Cat# 5221SC)

### CRISPR construct

The custom CRISPR gene editing constructs were obtained from Genecopoeia as an all-in-one CRISPR clone with a single guide RNA (sgRNA) targeting all variants for *CACNA1E* (target Site: CCTCAGGATGGCTCGCTTCG), *NTSR2* (target site: CCGCGCTCTACGCACTCATC), *TRPV1* (target site: CCCGGTGGATTGCCCTCACG) and the Cas9 gene, along with a neomycin resistance gene for mammalian cell selection and either an mCherry or GFP gene for visualization. (no. HCP001992-CG04-1B, HCP206276-CG04-1-B, HCP002057-CG12-1-B respectively). After the Cas9 sequence the plasmid carrying an mCherry reporter tag expresses downstream from an EF1a promoter while the GFP tag expresses downstream from T2A spacer sequence (**Fig. 2B**). Also included in the plasmids is a neomycin resistance gene for mammalian cell selection. Each DNA vector was amplified for use using standard bacterial transformation and an endotoxin-free maxi-prep Qiagen kit.

### Western blotting and analysis

Human DRG protein lysates were collected in ice cold lysis buffer (20 mM Tris-HCl (pH 7.4), 50 mM NaCl, 2 mM MgCl_2_ hexahydrate, 1% (*v*/*v*) NP40, 0.5% (*w*/*v*) sodium deoxycholate, 0.1% (*w*/*v*) SDS supplemented with protease and phosphatase inhibitor cocktail (BiMake)). The tissue lysate was centrifuged at 12,000× *g* for 10 min at 4 °C, then the supernatant was collected and used for determination of protein content with a BCA assay. Total protein (20-40 μg) from tissue supernatant was loaded into TGX precast gels (4–20% CriterionTM, BioRad) and transferred to nitrocellulose membrane (AmershamTM ProtranTM, GE Healthcare). After transfer, the membrane was blocked at room temperature for 1 h in blocking buffer (5% dry milk in Tris-buffered saline with tween 20 (TBST)). The membrane was incubated in diluted primary antibodies for 24h at 4 °C. The membrane was washed three times in TBST for 5 min each then incubated with peroxidase-conjugated secondary antibodies for one hour rocking at room temperature. The membrane was then washed in TBST for 5 min each, and then incubated with BioRad clarity western ECL substrate and Millipore HRP substrate solutions and then imaged in a ChemiDoc MP from BioRad. Some blots were then stripped with 25 mM glycine-HCl and 1% SDS (pH 2.0) for 30 to 60 min of rocking at room temperature before being washed and re-exposed to primary antibody. The resulting image bands were quantified using Scion Image (based on NIH Image). All images were quantified in the linear signal range. The intensities were normalized to GAPDH for Cav2.3, and for NTSR2 and TRPV1, the data was normalized to total protein. The normalized intensities were further normalized to a negative CRISPR control present on the same blot.

### Immunocytochemistry

The treated cultures were immediately fixed with 10% formalin (ThermoFisher Scientific, Cat# 23-245684) for 15 min at room temperature and rinsed three times with 1X PBS. 1 h block was performed at room temperature with 10% NGS and 0.3% Triton X-100 in 1X PBS. The coverslips were then incubated overnight at 4°C with primary antibodies anti-peripherin (chicken 1:1000; EnCor, AB_2284443) Cav2.3 (rabbit, Alomone labs, 1:100 AB_2039777; lot ACC006AN0302), NTSR2 (rabbit, 1:200, Thermo Fisher BS-12004R), TRPV1 (Rabbit, 1:500 Fisher scientific PA1748). Coverslips were rinsed 3 times with 1X PBS for 15 min and incubated with secondary antibodies Alexa Fluor goat anti-chicken IgG 647 (1:5000; Invitrogen A-21449, lot 2186435), Alexa Fluor goat anti-rabbit IgG 488 (1:5000; Invitrogen A-11034, lot 2110499) and Alexa Fluor goat anti-mouse IgG 594 (1:5000; Invitrogen, A11032, lot 1985396) for 1h at room temperature. The coverslips were washed 3 times with 1X PBS and mounted with prolong gold onto uncharged glass slides and allowed to cure overnight. Slides were imaged at 10X and 20X magnification using the Olympus FV1200 RS confocal laser scanning microscope.

### CRISPR insertion or deletion detection system with T7 Endonuclease I

36-48 hours post transfection of CRISPR plasmid constructs into cultured primary Human DRG the 6-well plate was removed from the incubator, media removed, and cells rinsed with 1 mL of PBS twice. Whole cell lysate preparation of primary cultured Human Dorsal Root Ganglia was collected by adding 200-300 uL of lysis buffer, provided in IndelCheck™ kit purchased from GeneCopoeia (Cat# IC001), into each well followed by use of a cell scrapper. Lysates were briefly vortexed before heating samples at 65°C for 20min, then 95°C for 10 min in a heat block. Lysates were quickly placed in ice for 1 minute before centrifugation at 12,000 RPM for 1 minute, supernatant of each sample was collected and placed in separate tube. Samples were then subjected to standard PCR. The 2x superhero PCR mix provided in the IndelCheck kit was used with primers purchased from GeneCopoeia specific to each target. Target site PCR primer for indel detection for NTSR2; Cat# HCP206276-CG04-1-B (NM_012344.3) (forward sequence: 5’- CGGTCCCACGTTGGCTCAGG -3’ and reverse sequence: 5’- ACCAGCACCGTGCACACTCG -3’); Target site PCR primer for indel detection for CaV2.3; Cat# HCP001992-CG04-1-B (Gene ID: 777), (forward sequence: 5’- CTGGGCACCCCCAAGCCCTA -3’; reverse sequence: 5’- CCCAAACGGGTCTCCACGGC -3’) and TRPV1; Cat# PHCP002057-a (Gene ID: 7442), (forward sequence: 5’- TTTCCTTGTTCTGTGTGGGG -3; reverse sequence: 5’- GACATTTAGCCCAGAAGCCA -3’). 8 uL of lysate was mixed with 1.25 uL of forward and reverse primers (5uM each) 12.5 uL 2xsuperhero PCR mix and 3.25 uL of nuclease free water for a total of 25 uL per sample. Samples placed into PCR tubes and put into Biorad thermocycler using the following settings: 94°C 5 minutes for 1 cycle; followed by 30 cycles at 94°C 30 seconds; 58°C for 30 seconds; 72°C for 1 minute; and finally, 1 cycle at 72°C for 5 minutes. For the denaturing and annealing process, 10 uL of each sample PCR product was mixed with 9 uL of nuclease free water for a total volume of 19 uL. Each mixture was mixed and centrifugated in a benchtop mini centrifuge for 30 seconds. Samples were placed in heat block set at 95°C for 5 minutes followed by a reannealing period at room temperature for 5 – 10 minutes. 1 uL of T7 endonuclease provided in kit was added to each sample and incubated at 37°C for 45-60 minutes. Samples were prepared with loading buffer and gel green to be loaded into wells of a 1.5% agarose DNA gel along with DNA ladder. Gel was run at 90V in TBE. Afterward, the gel was imaged using Biorad ChemiDoc Touch Imaging System imager.

### Fluorescent imaging plate reader-based calcium assay

Human dorsal root ganglion (hDRG) was collected from donor and subjected to above listed primary culture protocols and seeded into a clear flat-bottom black 96-well culture plate (Corning Inc 3603) coated with 100μg/mL PDL. The cells were maintained at 37°C, 5% CO2 for 24 hours. After 24 hours the cells were transfected with CRISPR/Cas-9 construct using lipofectamine 3000 and were left in the incubator for 18-24 hours at 37°C, 5% CO2 (**Table 2**). The media containing lipid/DNA complex was replaced with regular hDRG media after 18–24-hour incubation. 48-hours after media replacement the plate was pulled from the incubator and media removed from the wells, each well was washed once with 200uL of Tyrode’s buffer (NaCl 119 mM, KCl 2.5 mM, MgCl2 2 mM, CaCl2 2 mM, HEPES 25 mM, Glucose 30 mM). Following wash, 150uL of Tyrode’s containing 2 uM Fura 2-AM (Invitrogen F1221, Lot 2559176)/ Pluronic F-127 (Invitrogen P3000MP, lot 2510678) (mixed 1:1 by volume) was added to each well containing cells. The plate was incubated while covered at room temperature for an hour. The dye solution was removed, the wells were washed twice with Tyrode’s buffer before adding 150uL of Tyrode’s buffer supplemented with 2.5 mM probenecid (Invitrogen P36400, lot 2600149), left to rest covered at room temperature for 20-30 minutes, followed by an extra wash with Tyrode’s buffer supplement with probenecid. After incubation period, the plate was placed into Tecan plate reading instrument for five time point baseline. Plate was removed and media replaced. The NC and KO wells received 150 uL of Tyrode’s supplemented with probenecid with 800 nM capsaicin (Sigma M2028-50MG, Lot SLCB0726) while the non-stimulated control well received only Tyrode’s supplemented with probenecid. The plate was placed back into the instrument and 12 points were read post capsaicin treatment. Each read point was 30 seconds apart. See **table 3** for instrument reader parameters.

**Table 3.**
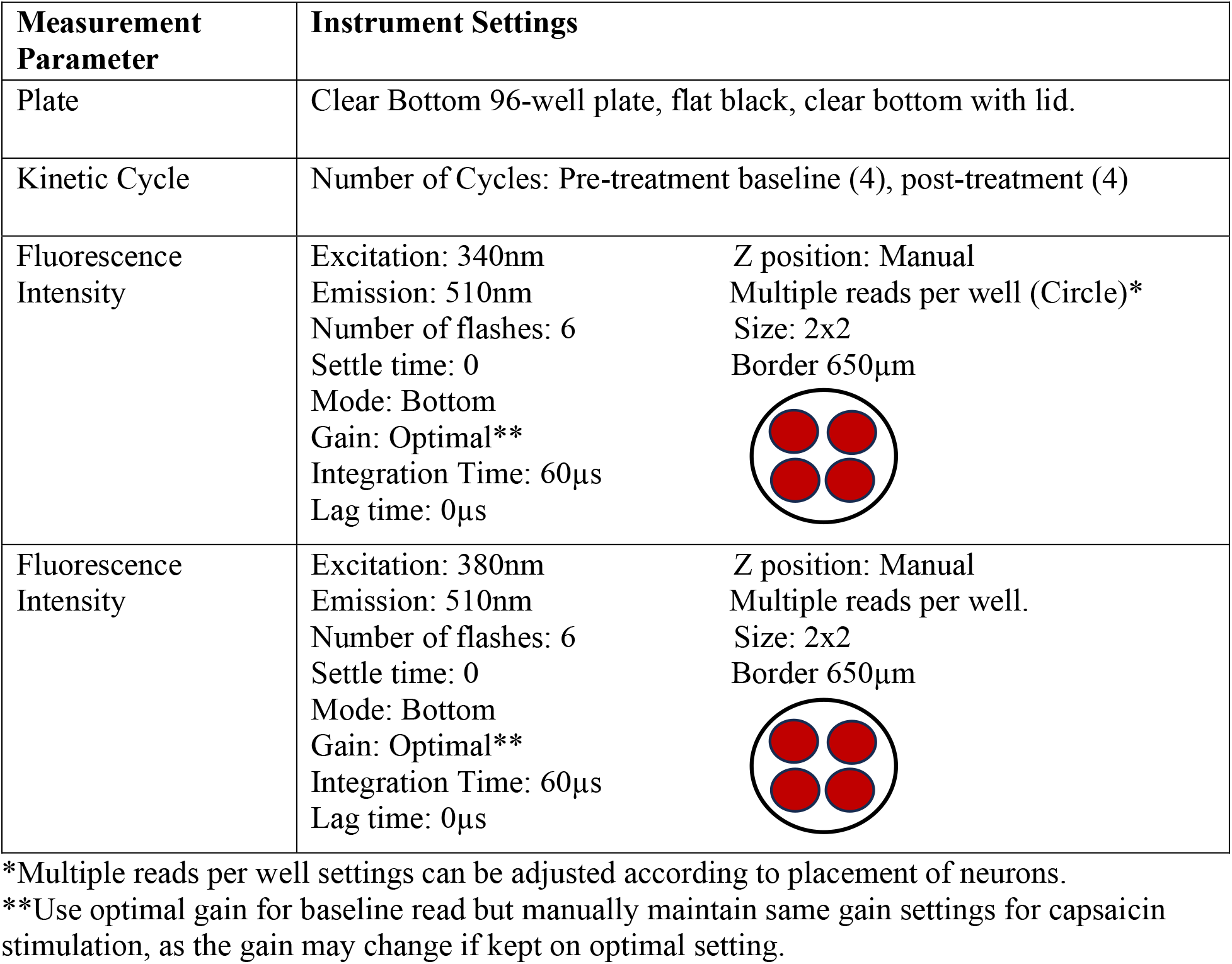
FLPR instrument settings.

| Measurement Parameter | Instrument Settings |  |
| --- | --- | --- |
| Plate | Clear Bottom 96-well plate, flat black, clear bottom with lid. |  |
| Kinetic Cycle | Number of Cycles: Pre-treatment baseline (4), post-treatment (4) |  |
| Fluorescence Intensity | Excitation: 340nm<br>Emission: 510nm<br>Number of flashes: 6<br>Settle time: 0<br>Mode: Bottom<br>Gain: Optimal**<br>Integration Time: 60μs<br>Lag time: 0μs | Z position: Manual<br>Multiple reads per well (Circle)*<br>Size: 2x2<br>Border 650μm |
| Fluorescence Intensity | Excitation: 380nm<br>Emission: 510nm<br>Number of flashes: 6<br>Settle time: 0<br>Mode: Bottom<br>Gain: Optimal**<br>Integration Time: 60μs<br>Lag time: 0μs | Z position: Manual<br>Multiple reads per well.<br>Size: 2x2<br>Border 650μm |

\*Multiple reads per well settings can be adjusted according to placement of neurons.
\*\*Use optimal gain for baseline read but manually maintain same gain settings for capsaicin stimulation, as the gain may change if kept on optimal setting.

### Viability Assays

Two different viability dyes were utilized in the experiments: CellTiter Glo 2.0 and the CellTiter 96 Aqueaous one solution cell proliferation assay (MTS). For both assays we chose to use a 96-well plate and seeded the hDRG cultures at a density of at least 150 neurons per well. For the MTS assay, we preincubated 20ul of the dye per 100ul of media for 2 hours at 37°C, 5% CO2, and recorded the absorbance at 490nm. Media was completely changed, and the cells were either transfected with the *TRPV1* gRNA plasmid, lipofectamine without plasmid, or just media for 24 hours at 37°C, 5% CO2. Media was changed and 20ul of the MTS dye was added to the cells per 100ul of media and absorbance was recorded two hours later at 490nm. The percent change in viability was determined using the following equation ((post-treatment MTS – Pre-treatment MTS)/Average Pre-treatment MTS)) *100 (*43*). For the CellTiter Glo 2.0, we added 1:1 ratio of media to dye to the wells post treatment with either the *TRPV1, CACNA1E, NTSR2* gRNA plasmid (500ng), lipofectamine alone, or media alone. The plate was placed in an orbital shaker at room temp for 2 minutes to induce cell lysis and luminescence was recorded 10 minutes later. Both assays were recorded using a Tecan Spark 20 plate reader.

